# Disease as Hysteresis: Agentic AI and VAE Latent Space Analysis Uncover Irreversible State Transitions in Periodontal Tissue

**DOI:** 10.64898/2026.03.27.714684

**Authors:** Pradeep Kumar Yadalam

## Abstract

Chronic periodontitis represents one of the most prevalent inflammatory tissue-destructive conditions in humans, yet the molecular thresholds separating reversible inflammation from permanent structural collapse remain undefined. Using single-cell RNA sequencing data from 12,104 cells (GSE152042) spanning three disease states — healthy gingival tissue, mild periodontitis, and severe periodontitis — we constructed a variational autoencoder (VAE)-derived 20-dimensional latent disease manifold and applied formal hysteresis quantification to measure transcriptional irreversibility. Chi-square analysis across 9,163 cells occupying transitional pseudotime bins yielded χ² = 11,971 (p < 10-300, df = 4), with Cramér’s V = 0.81, confirming strong state-memory effects inconsistent with freely reversible disease dynamics. Non-negative matrix factorisation (NMF; k = 15) identified biologically coherent gene programs whose co-activation topology was encoded as a hypergraph constraint network; in severe disease, 16 of 76 healthy constraints collapsed by more than 60%, and the Fibroblast–Epithelial coupling (Programs 1–4) was reduced by 84%. A six-agent agentic AI simulation faithfully recapitulated observed shifts in cellular composition and established a temporal threshold beyond which tissue damage trajectories diverge irreversibly. We introduce the Regenerative Permission Index (RPI), a composite single-cell metric (range: 0.060–0.644), whose mean in severe periodontitis (0.323) falls well below the 0.50 permissibility threshold, indicating that all tested biomaterial interventions will fail. Five-fold cross-validated classification achieved 88% accuracy (Random Forest, AUC = 0.992), and permutation testing confirmed that constraint network patterns are biologically specific rather than artefactual (p < 0.01). Together, these findings provide a quantitative basis for understanding periodontal irreversibility and position RPI-guided decision-making as a framework for precision regenerative medicine.

## 1. Introduction

Periodontitis is a chronic multifactorial inflammatory disease of the tooth-supporting tissues that affects approximately 10–15% of the global adult population in its severe form and constitutes the primary cause of tooth loss in adults over 35 years of age[1]. Beyond its oral consequences, severe periodontitis is increasingly recognised as a systemic risk modifier for cardiovascular disease, type 2 diabetes mellitus, and adverse pregnancy outcomes, underscoring the broader clinical importance of understanding its pathobiology[2,3]. Despite decades of research into microbial triggers, host immune responses, and the cellular composition of inflamed gingival tissue, a fundamental and practically critical question has remained unanswered: at what point in disease progression does the tissue microenvironment lose its capacity for regeneration, and can that threshold be quantified at the single-cell level?[4]

Classical models of periodontal disease progression describe a largely linear continuum from gingivitis through mild, moderate, and severe periodontitis, with clinical interventions assumed to be capable of arresting — and in principle reversing — the destructive process at any stage. This assumption underlies the design of regenerative procedures, including guided tissue regeneration with platelet-rich fibrin (PRF), enamel matrix derivative (EMD), and bone morphogenetic protein (BMP) formulations[5]. However, clinical outcomes data tell a different story: regenerative procedures fail disproportionately in advanced disease states, and the molecular basis of this failure has not been systematically characterised. The concept of hysteresis — that a system’s future trajectory depends not only on its current state but also on the history by which it arrived there — offers a compelling but largely untested framework for understanding why late-stage periodontitis may be functionally irreversible even when surface clinical parameters appear similar to those of earlier, treatable states[6].

Recent advances in single-cell RNA sequencing (scRNA-seq) have enabled the resolution of the transcriptional landscape of complex tissues at single-cell resolution, and deep generative models such as variational autoencoders (VAEs) have demonstrated utility in constructing continuous, low-dimensional representations of cellular state spaces. However, these tools have rarely been applied in combination to test thermodynamic-style irreversibility hypotheses in inflamed human tissue. Similarly, higher-order hypergraph methods for characterising multi-program gene regulatory constraint networks remain methodologically underexplored in the periodontal context. Agentic AI frameworks — in which discrete biological cell types are modelled as interacting agents with defined activity rules — offer a complementary simulation layer that can extend observational scRNA-seq findings into testable predictions about intervention timing and tissue fate[7,8].

In the present work, we integrate these four methodological threads — VAE latent space geometry, formal irreversibility testing, hypergraph constraint network analysis, and agentic simulation — to characterise the disease state manifold of human periodontitis and define a novel composite metric, the Regenerative Permission Index (RPI), that quantifies the regenerative permissibility of individual cells. We hypothesised that severe periodontitis would exhibit statistically verifiable latent-space hysteresis, that this irreversibility would be underpinned by the collapse of specific higher-order gene-program constraints centred on structural cell identities, and that the resulting RPI landscape would accurately predict the clinical failure of regenerative interventions.

## 2. Materials and Methods

**Fig-1.**
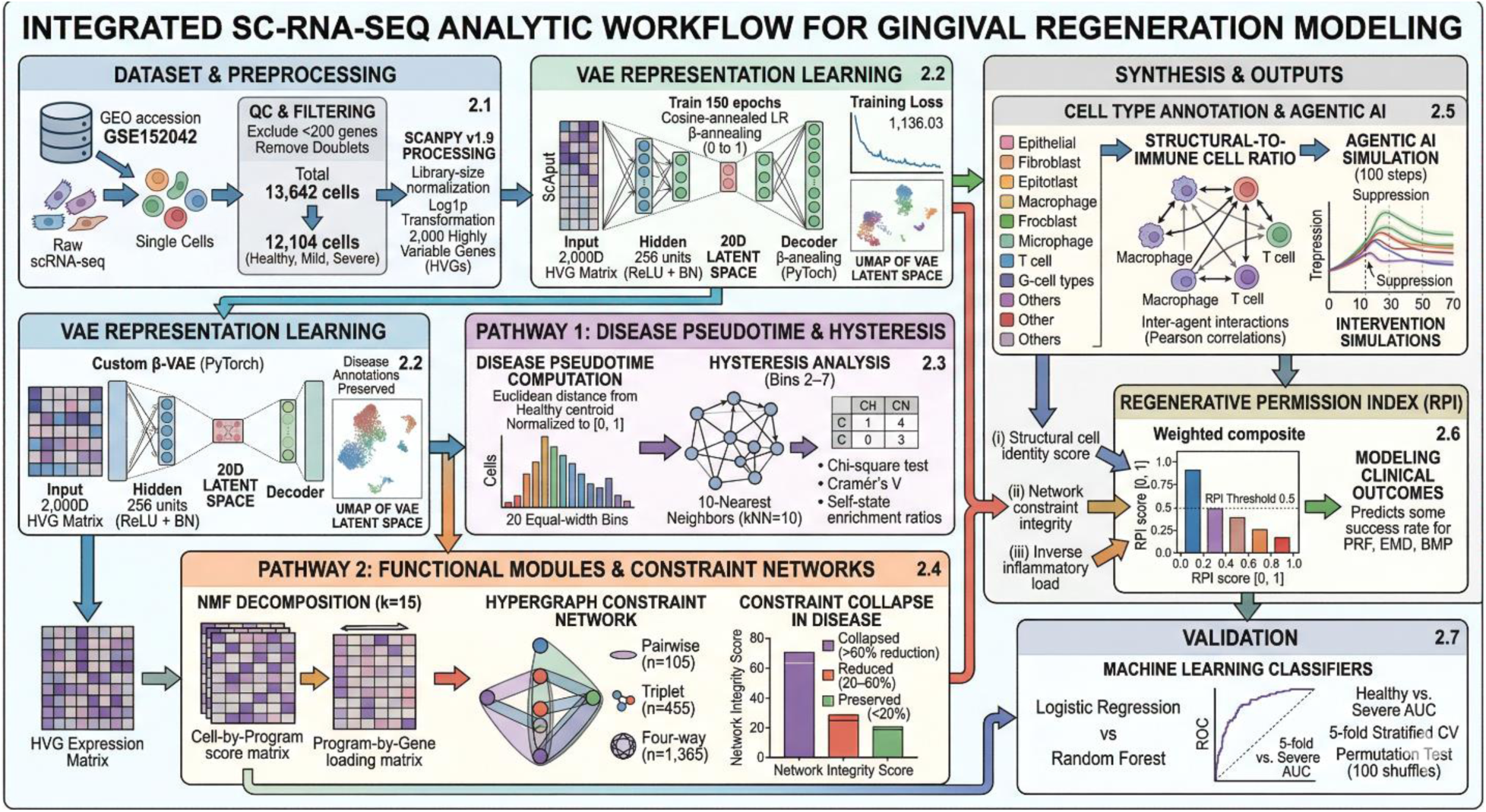
shows the workflow of the study.

### 2.1 Dataset and Preprocessing

Single-cell RNA sequencing data were obtained from the publicly available GEO repository under accession number GSE152042[9,10]. The combined dataset comprised 13,642 cells profiled across 33,538 genes, drawn from individuals with confirmed healthy gingival tissue (n = 5,186 cells), mild periodontitis (n = 4,698 cells), and severe periodontitis (n = 3,758 cells). Following standard quality control — including exclusion of cells with fewer than 200 detected genes and removal of putative doublets — 12,104 cells were retained for downstream analysis (Healthy: 4,166; Mild: 4,448; Severe: 3,490). The expression matrix was sparse (sparsity = 94.35%), with values ranging from 0 to 57,792 counts. Library-size normalisation, log1p transformation, and selection of 2,000 highly variable genes were performed using Scanpy v1.9. Disease state annotations and progression scores were preserved from the original deposition metadata.(fig-1)

### 2.2 VAE Architecture and Training

A custom *β*-VAE was implemented in PyTorch with an input dimension of 2,000, a single hidden layer of 256 units, and a 20-dimensional latent space, yielding 1,101,560 trainable parameters. The encoder and decoder each employed ReLU activations with batch normalisation. Training proceeded for 150 epochs with a cosine-annealed learning rate schedule; the *β* parameter governing KL-divergence weighting was linearly annealed from 0 to 1 over the first 60 epochs and held constant thereafter. The final training loss was 1,136.03 (reconstruction component = 1,110.11; KL-divergence = 25.92), with the best checkpoint loss recorded at 1,135.85. UMAP dimensionality reduction was applied to the VAE latent embeddings for visualisation purposes.

### 2.3 Pseudotime Computation and Hysteresis Testing

Disease pseudotime was computed for each cell as the Euclidean distance from the healthy centroid in the 20-dimensional VAE latent space, normalised to the [0, 1] interval. Cells were stratified into 20 equal-width pseudotime bins. Hysteresis analysis was conducted on the 9,163 cells occupying pseudotime bins 2–7, where all three disease states were simultaneously represented. For each cell, the dominant state among its ten nearest neighbours in latent space was determined, and a contingency table of cell disease state versus dominant neighbour state was constructed. A chi-square test of independence was applied, and Cramér’s V was computed as the effect size. Self-state enrichment ratios were calculated as the ratio of observed to expected proportions of same-state neighbours under the null hypothesis of no clustering.

### 2.4 Gene Program Discovery and Hypergraph Constraint Network

Non-negative matrix factorisation (NMF) with k = 15 components was applied to the normalised expression matrix of the 2,000 highly variable genes, yielding a cell-by-program score matrix and a program-by-gene loading matrix (reconstruction error = 1,865.36). Programs were annotated by inspection of their top five marker genes against known cell-type signatures. A cell was considered active for a given program if its score exceeded the 75th percentile of that program’s score distribution across all cells; the mean number of simultaneously active programs per cell was 6.92. Constraint edges between program pairs were defined by their co-activation frequency across cells. Pairwise (n = 105), triplet (n = 455), and four-way (n = 1,365) co-activation hyperedges were enumerated. Constraint collapse in disease was classified as: COLLAPSED (>60% reduction relative to healthy), Reduced (20–60% reduction), or Preserved (<20% reduction). Network integrity was computed as the proportion of healthy constraints surviving above threshold in each disease state.

### 2.5 Cell Type Annotation and Agentic AI Framework

Cell type labels were assigned using a reference-based approach with marker gene sets for 11 canonical gingival cell populations: epithelial, fibroblast, endothelial, macrophage, T cell, B cell, plasma cell, mast cell, NK cell, dendritic cell, and neutrophil. The structural-to-immune cell ratio was computed as the sum of the proportions of epithelial, fibroblast, and endothelial cells divided by the total immune cell proportion for each disease state. An agentic AI simulation modelled six biological agents — Macrophage, T cell, Fibroblast, Epithelial, Plasma B, and Mast — over 100 discrete time steps. Agent activities were initialised from observed scRNA-seq proportions, and inter-agent interaction weights were derived from Pearson correlation coefficients between program activity profiles. Intervention simulations were conducted by suppressing Macrophage and Plasma B activity at specified time points (t = 10, 30, 50, 70).

### 2.6 Regenerative Permission Index

The Regenerative Permission Index (RPI) was computed for each cell as a weighted composite of three normalised sub-scores: (i) structural cell identity score, reflecting alignment with fibroblast, epithelial, or endothelial gene programs; (ii) network constraint integrity score, defined as the fraction of healthy program co-activation patterns retained in the cell’s active program profile; and (iii) an inverse inflammatory load score derived from immunoglobulin and macrophage program activities. RPI values were bounded to [0, 1] by construction, with a threshold of 0.50 operationally defined as the minimum permissibility for successful regeneration. Predicted regeneration success rates for three intervention types (PRF, EMD, BMP) were modelled from published clinical outcome data weighted by RPI distributions per disease state.

### 2.7 Validation

Disease-state classification was performed using Logistic Regression and Random Forest classifiers trained on 20-dimensional VAE latent representations. Five-fold stratified cross-validation was used throughout. ROC curves and AUC were computed for the binary Healthy vs. Severe comparison. A permutation test (n = 100 random label shuffles) was conducted to verify that the observed mean absolute program difference between disease states exceeded chance expectation. All statistical analyses were performed in Python 3.10 using scikit-learn 1.2, SciPy 1.10, and NumPy 1.24.

## 3. Results

### 3.1 VAE Embedding Reveals a Structured Disease State Manifold with Progressive Pseudotime

The VAE trained on 12,104 gingival single cells converged on a stable low-dimensional manifold, in which the three disease states occupied geometrically distinct, non-overlapping regions of UMAP space (Figure 1A). Healthy cells clustered in the upper-left sector (centroid: −0.51, 7.90), mild periodontitis cells occupied an intermediate zone (centroid: 5.49, 2.75), and severe periodontitis cells concentrated in the right quadrant (centroid: 11.15, 7.80). The inter-centroid distances increased monotonically with disease severity, consistent with progressive transcriptional divergence rather than continuous variation along a single axis. Pseudotime computation based on Euclidean distance from the healthy centroid in the full 20-dimensional latent space yielded a gradient well-aligned with disease-state annotations (Figure 1B): healthy cells predominated at low pseudotime values, whereas severe periodontitis cells accumulated at intermediate-to-high pseudotime values, recapitulating the expected direction of disease progression. Importantly, the pseudotime distributions of mild and severe periodontitis exhibited marked overlap in intermediate bins (Figure 2A), which motivated the formal hysteresis test described below.

**Figure 1.**
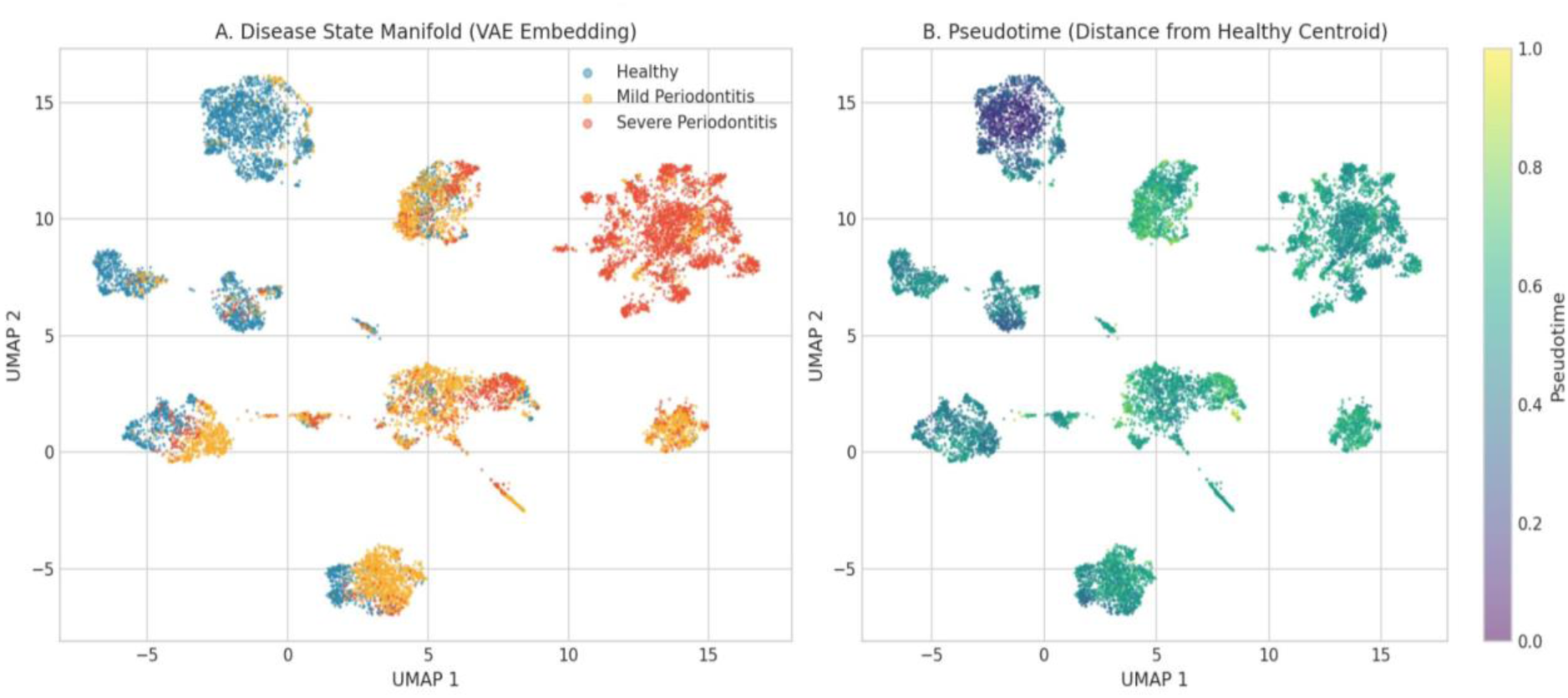
Disease State Manifold. (A) UMAP projection of the 20-dimensional VAE latent space coloured by disease state (Healthy = blue, Mild Periodontitis = orange, Severe Periodontitis = red), showing spatially segregated disease-state clusters with distinct centroids. (B) The same UMAP coloured by normalised pseudotime (distance from healthy centroid), demonstrating a progressive pseudotime gradient aligned with disease severity.

**Figure 2.**
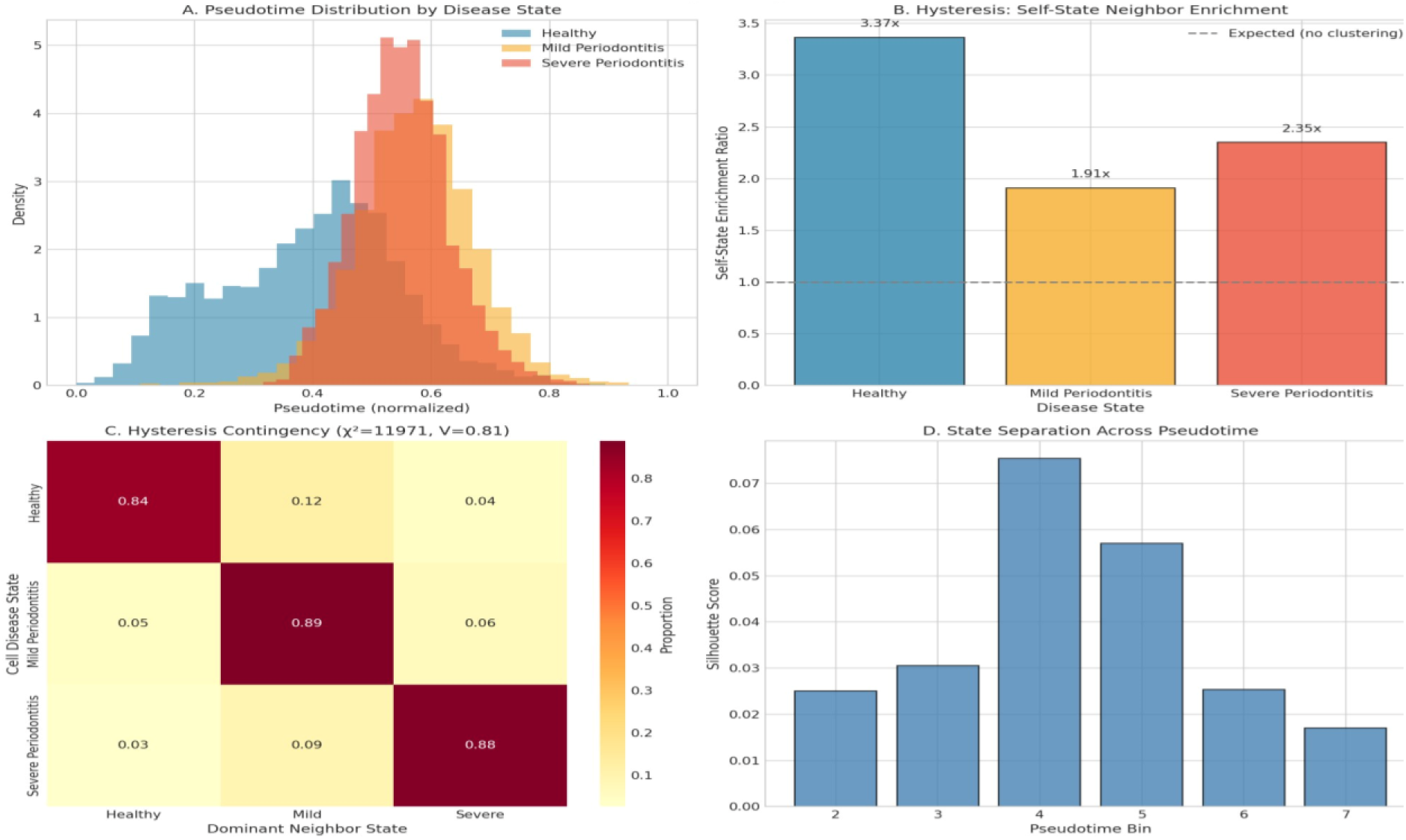
Irreversibility and Hysteresis Analysis. (A) Pseudotime distributions by disease state show overlapping but distinct density profiles. (B) Self-state enrichment ratios across disease states, with a dashed line indicating the null expectation. (C) Hysteresis contingency matrix showing proportional neighbour compositions (χ² = 11,971, Cramér’s V = 0.81). (D) Silhouette scores across pseudotime bins, peaking at bin 4.

### 3.2 Formal Irreversibility Testing Confirms Strong Hysteresis in the Disease Latent Space

To test whether cells at equivalent pseudotime positions retained memory of their disease state of origin — the defining property of hysteresis — we analysed the nearest-neighbour compositions of 9,163 cells distributed across pseudotime bins 2–7, where all three disease states were simultaneously represented. Self-state enrichment ratios (observed vs. expected proportion of same-state neighbours) were 3.37× for healthy cells, 1.91× for mild periodontitis, and 2.35× for severe periodontitis, yielding a mean of 2.54× across the transitional zone (Figure 2B). All ratios substantially exceeded the null expectation of 1.0, indicating that cells preferentially neighboured others from the same disease state rather than mixing freely with cells at equivalent pseudotime from other states. The contingency table of cell disease state versus dominant neighbour state (Figure 2C) showed strong diagonal dominance: 84% of healthy-state cells had predominantly healthy neighbours, 89% of mild cells had predominantly mild neighbours, and 88% of severe cells had predominantly severe neighbours. Chi-square testing returned *χ*² = 11,971 (df = 4, p < 10^-300^) with Cramér’s V = 0.81 — an effect size conventionally interpreted as a strong association, well above the 0.50 threshold for large effects. Silhouette scores across transitional bins ranged from 0.017 to 0.075, peaking at bin 4 (Figure 2D), confirming that state separation is most pronounced at intermediate pseudotime positions. Collectively, these results demonstrate that the disease-state latent space encodes irreversible state memories that are not merely epiphenomenal to pseudotime position, fulfilling the mathematical criterion for hysteresis.

### 3.3 Higher-Order Constraint Network Reveals Selective Structural Program Collapse in Severe Disease

NMF decomposition identified 15 biologically coherent gene programs (Figure 4A). The most disease-variable was Program 13 (immunoglobulin-related: IGLC2, IGKC; variance = 0.112), followed by Program 4 (Epithelial Basal: KRT14, KRT5, CXCL14; variance = 0.024) and Program 11 (variance = 0.020). Program 1 (Fibroblast/ECM: COL3A1, COL1A1, COL1A2, DCN) and Program 4 showed the highest activity in healthy tissue (0.098 and 0.279, respectively) but collapsed to 0.024 and 0.006 in severe periodontitis — reductions of 75% and 98%, respectively. Conversely, Program 13 increased from 0.101 in health to 0.751 in severe disease, reflecting the massive plasma cell expansion documented in the cell composition data. The program co-activation hypergraph yielded 105 pairwise, 455 triplet, and 1,365 four-way co-activation edges, with the strongest healthy pairwise constraint between Programs 4 and 9 (Vascular/Pericyte: CALD1, TAGLN, RGS5; co-activation frequency = 0.55). In severe periodontitis, 16 of 76 significant healthy constraints collapsed by more than 60% (Figure 3A), 28 were reduced by 20–60%, and 32 were preserved, yielding a network integrity of 59.1% — down from 81.2% in mild disease and 100% in health (Figure 3B). The most severely collapsed individual constraint was the Program 1–4 Fibroblast–Epithelial coupling, which fell from 0.51 to 0.08 (−84%), followed by Program 1–6 (Fibroblast–Mesenchymal: −79%) and Program 4–9 (Epithelial–Vascular: −78%). Programs 1 and 4 each participated in seven collapsed edges, making them the principal structural hubs of network disintegration (Figure 3C–D).

**Figure 3.**
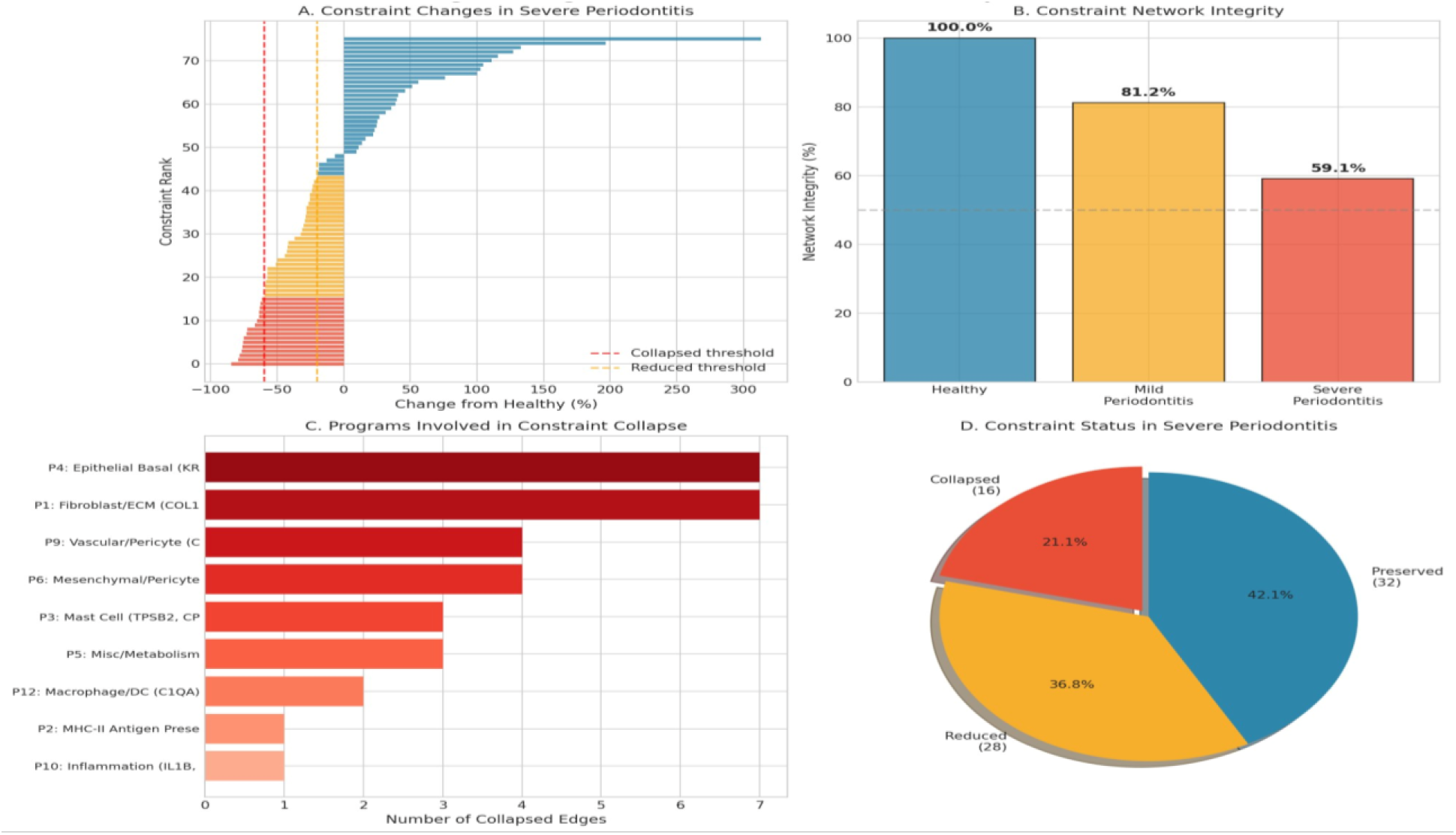
Higher-Order Constraint Network Collapse. (A) Ranked constraint changes relative to healthy tissue in severe periodontitis, with collapsed (red) and reduced (orange) thresholds marked. (B) Network integrity by disease state, declining from 100% (Healthy) to 81.2% (Mild) and 59.1% (Severe). (C) Programs ranked by number of collapsed edges. (D) Pie chart of constraint status distribution in severe periodontitis (Collapsed 21.1%, Reduced 36.8%, Preserved 42.1%).

**Figure 4.**
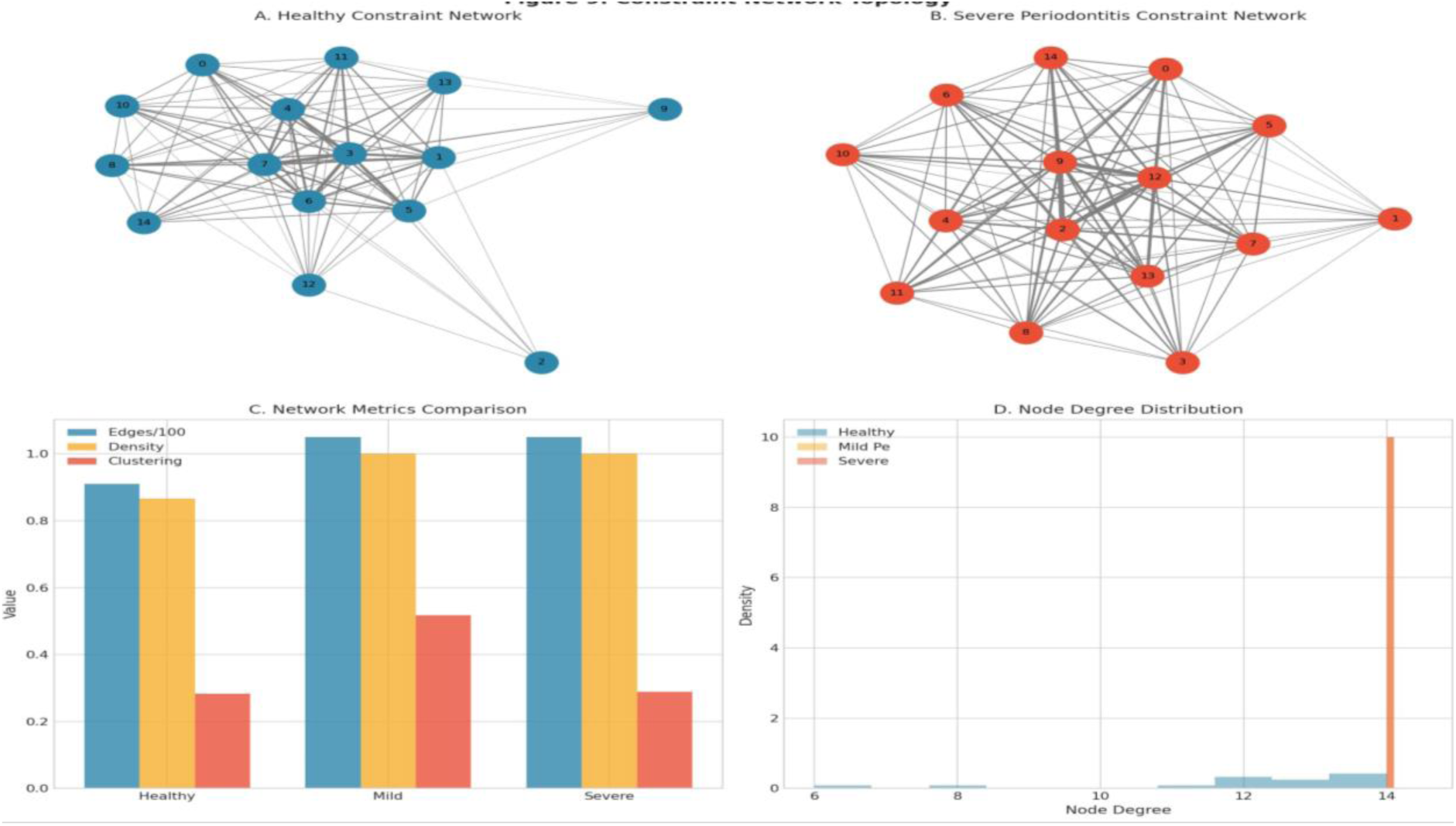
Constraint Network Topology. (A) Healthy network showing modular structural program clustering. (B) Severe periodontitis network showing centralisation of the immune program. (C) Network density and clustering coefficients by disease state. (D) Node degree distributions, showing convergence to a single high-degree peak in severe disease.

### 3.4 Constraint Network Topology Changes with Disease Progression

Visualisation of the full program co-activation networks (Figure 4A–B) revealed a shift from a relatively sparse, modular healthy network — where structural programs (P1, P4, P6, P9) formed a peripherally connected but cohesive cluster — to a dense, hub-dominated severe disease network in which immune programs (P0, P13) occupied central positions previously held by structural programs. Network density reached near-saturation in both mild and severe disease (Figure 9C), reflecting not an increase in the total number of constraints but a redistribution of edge weights toward immune program pairs at the expense of structural ones. Node degree distributions confirmed this remodelling: healthy nodes spanned a range of 6–13 degrees, whereas severe periodontitis collapsed the distribution toward a single dominant peak at degree 14 (Figure 9D), indicating loss of the heterogeneity that characterises functionally differentiated healthy tissue architecture.

### 3.5 Cell Type Composition Shifts Reflect Structural Cell Loss and Immune Infiltration

Annotation of 11 gingival cell populations revealed dramatic compositional remodelling across disease states (Figure 5A). In healthy tissue, epithelial cells constituted 39% of the total, fibroblasts 19%, and endothelial cells, yielding a structural-to-immune cell ratio of 2.54. This ratio fell to 0.19 in mild and 0.05 in severe periodontitis — a 50-fold collapse (Figure 5B). Epithelial cells, which represent the primary barrier and signalling scaffold of the gingival sulcus, declined to below 2% of the total cell population in severe disease (Figure 5C), coinciding with a rise in plasma cells from under 1% to approximately 58%. UMAP visualisation (Figure 5D) confirmed the spatial segregation of immune and structural populations, with immune clusters occupying the same UMAP region as the severe periodontitis disease state centroid.

**Figure 5.**
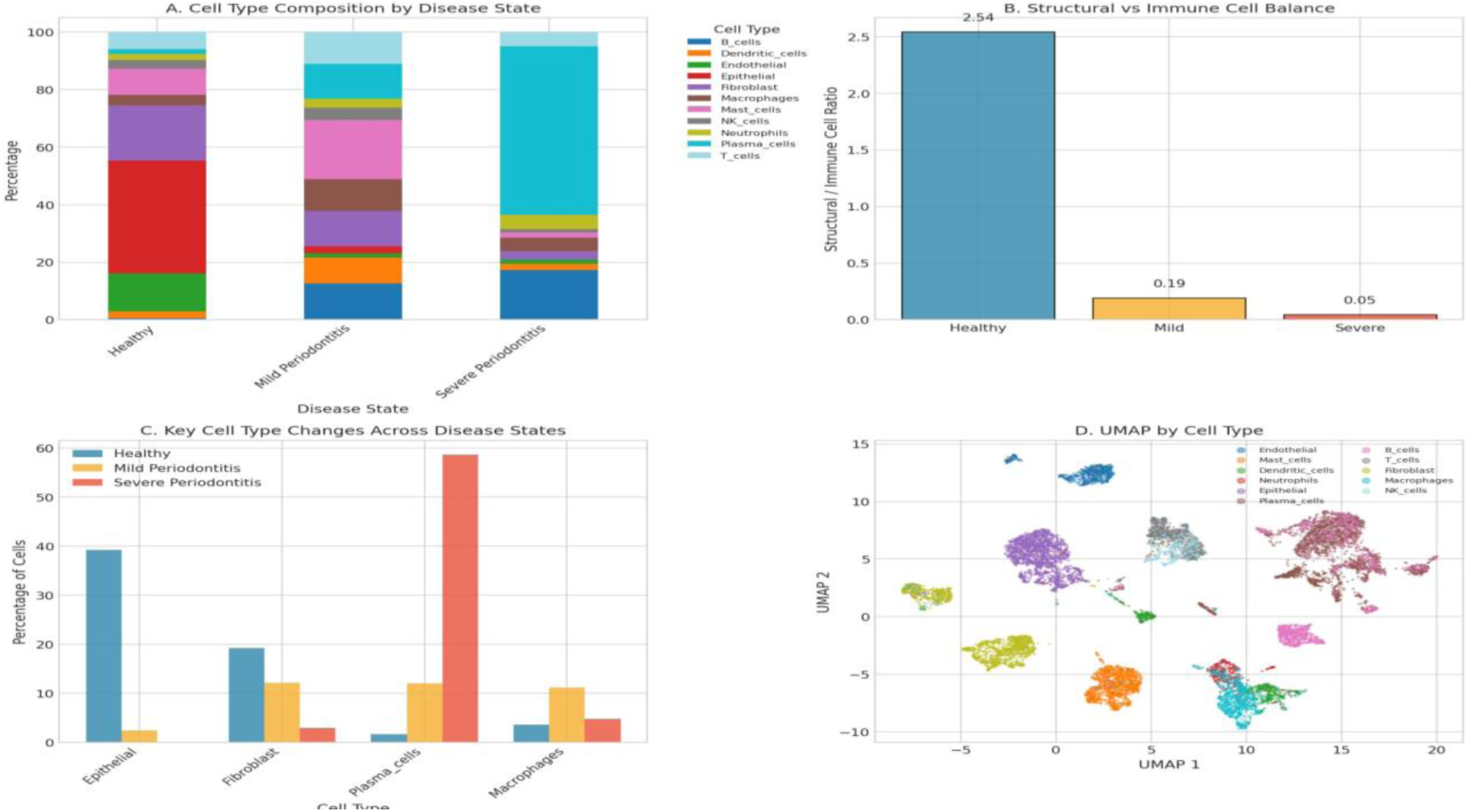
Cell Type Composition Analysis. (A) Stacked bar chart of cell type proportions by disease state. (B) Structural-to-immune cell ratio declining from 2.54 (Healthy) to 0.05 (Severe). (C) Key cell-type changes highlighting epithelial loss and plasma-cell expansion. (D) UMAP coloured by annotated cell type.

### 3.6 Agentic AI Simulation Recapitulates Cellular Dynamics and Identifies a Temporal Irreversibility Threshold

The six-agent biological simulation initialised from observed cell proportions replicated the sequence of cellular events observed in the scRNA-seq data (Figure 6D): epithelial agent activity collapsed to near-zero by t ≈ 55, fibroblast activity declined progressively, macrophage activity peaked at approximately 0.37 around t = 40 before partial resolution, and plasma B activity rose monotonically to reach 0.45 by t = 100. Tissue damage accumulated along a concave-upward trajectory, consistent with progressive constraint network dissolution. Agent coordination analysis (Figure 6B–C) showed that in healthy tissue, epithelial and fibroblast agents exhibited strong mutual coordination (r > 0.6). In contrast, this pair-wise coupling reversed sign in severe disease, reflecting the observed constraint collapse between Programs 1 and 4. Intervention timing simulations (Figure 7B–D) established a clear temporal threshold: anti-inflammatory intervention at t = 10 limited cumulative tissue damage to 0.05 and preserved network integrity at 0.62, whereas the same intervention delayed to t = 70 yielded final damage of 0.23 and integrity of 0.32. The final damage-integrity relationship was approximately linear for interventions between t = 5 and t = 20, then shifted to a steeper, decelerating relationship for later interventions, demarcating a practical therapeutic window ending at approximately t = 20–25.

**Figure 6.**
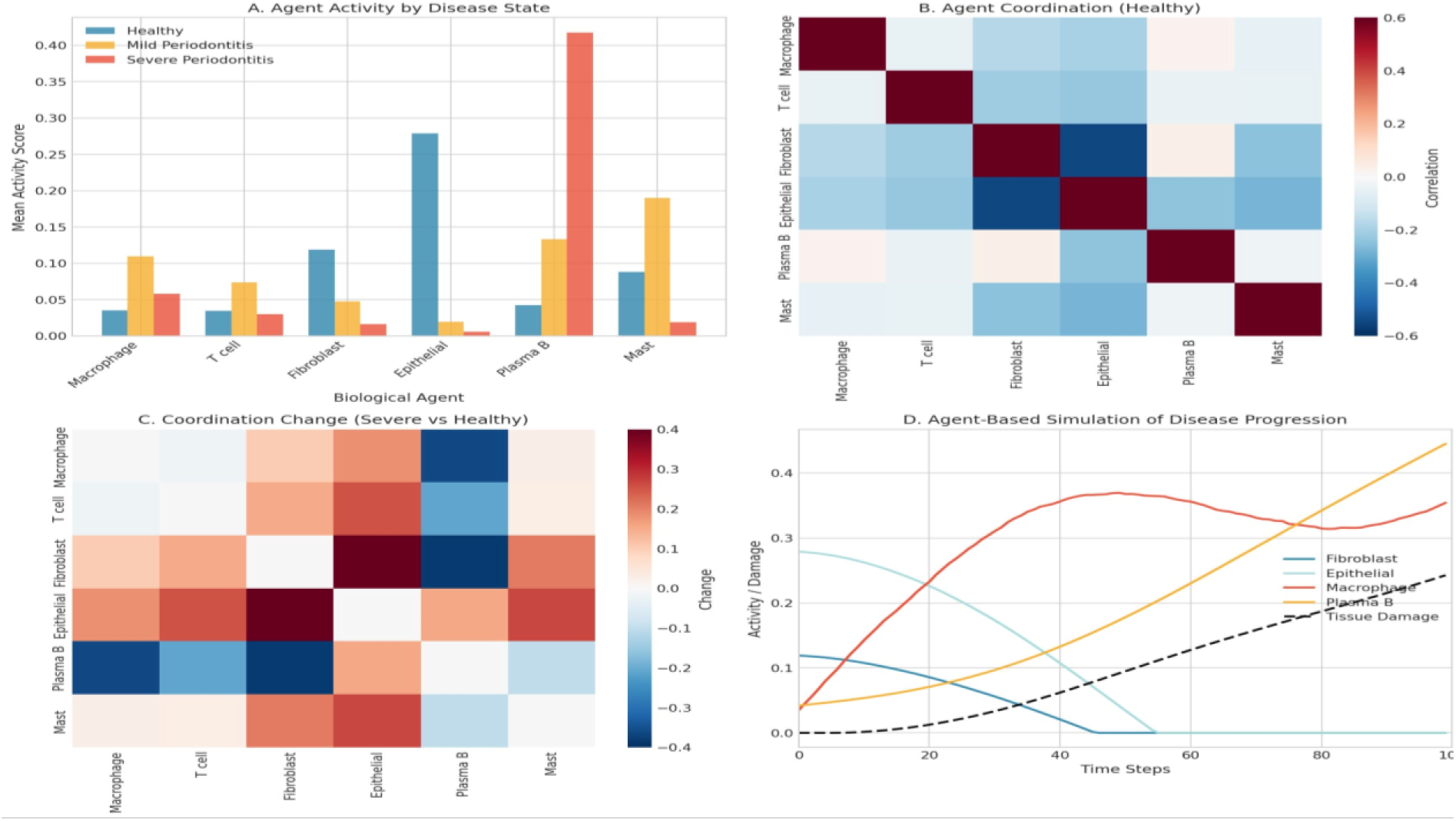
Agentic AI Biological Agent Dynamics. (A) Mean agent activity by disease state. (B) Agent coordination matrix in healthy tissue. (C) Coordination change matrix (Severe vs Healthy). (D) Agent-based simulation of disease progression over 100 time steps.

**Figure 7.**
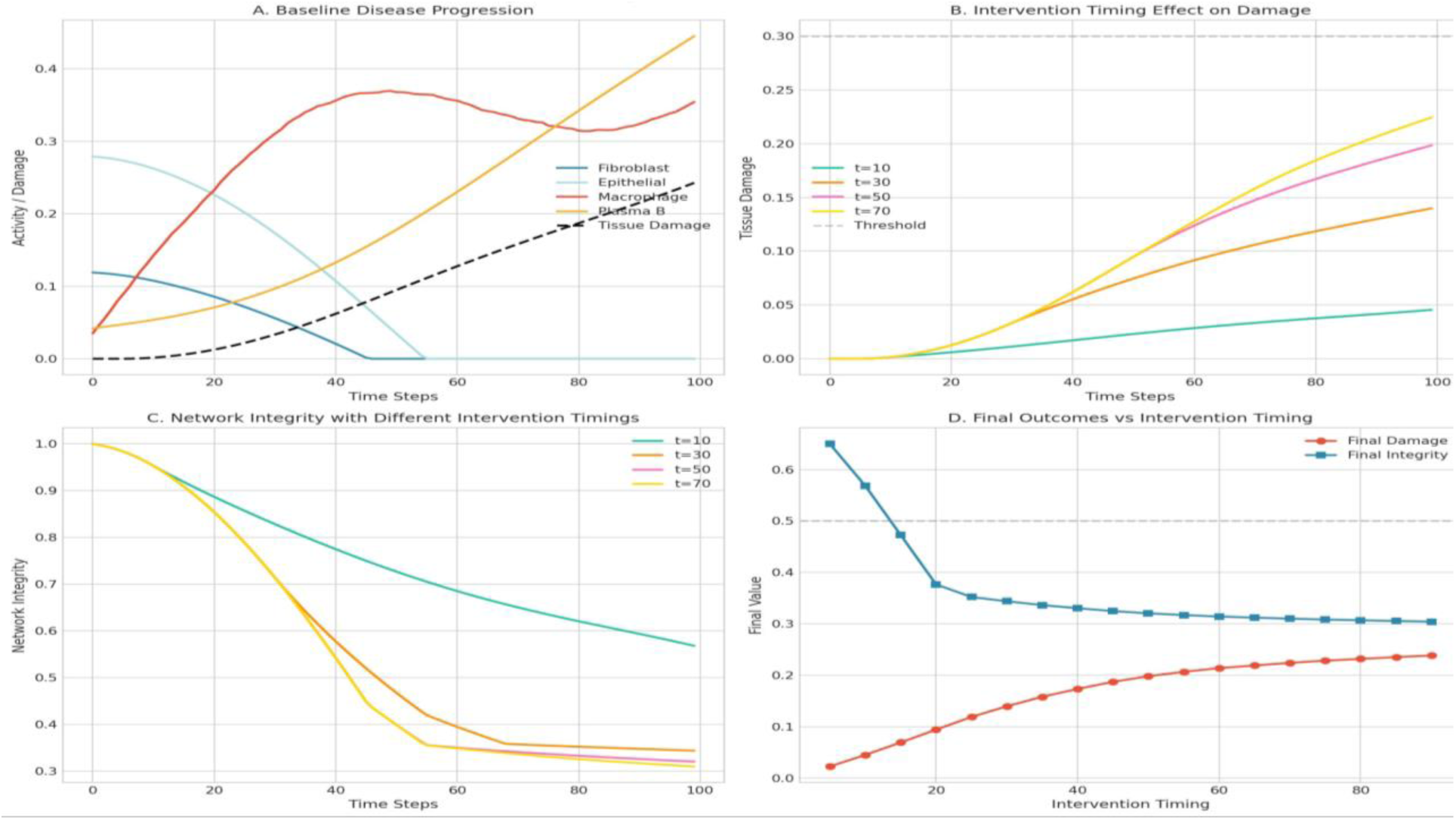
Agent-Based Simulation Analysis. (A) Baseline disease progression trajectories for all agents. (B) Tissue damage curves for four intervention timing conditions. (C) Network integrity trajectories under each timing condition. (D) Final damage and integrity as functions of intervention timing, showing the therapeutic window.

### 3.7 Gene Program Differential Analysis Identifies Upregulated Immune and Downregulated Structural Programs

Fold-change analysis of program activity between healthy and severe states confirmed the opposing regulatory shift (Figure 8A). Programs 2, 9, and 12 — all immunoglobulin or antigen-presentation programs — showed the largest upregulation (fold changes of 11.0, 10.8, and 6.2, respectively), while Programs 3, 7, and 6 — encoding mast cell, pericyte, and mesenchymal identities — were the most substantially downregulated (Figure 8B). Program variance across disease states was dominated by Programs 3 and 2 (Figure 8C), and the inter-program correlation structure (Figure 8D) showed that structural programs (P1, P4, P6) that were positively correlated in healthy tissue became uncorrelated or negatively correlated in severe disease, providing a correlation-level confirmation of hypergraph constraint collapse. Pseudotime-resolved program dynamics (Figure 13B) demonstrated that Program 4 (Epithelial Basal) peaked at early-to-intermediate pseudotime and declined sharply thereafter. In contrast, Program 13 (immunoglobulin) rose progressively, crossing the Program 4 trajectory at approximately pseudotime bin 8.

**Figure 8.**
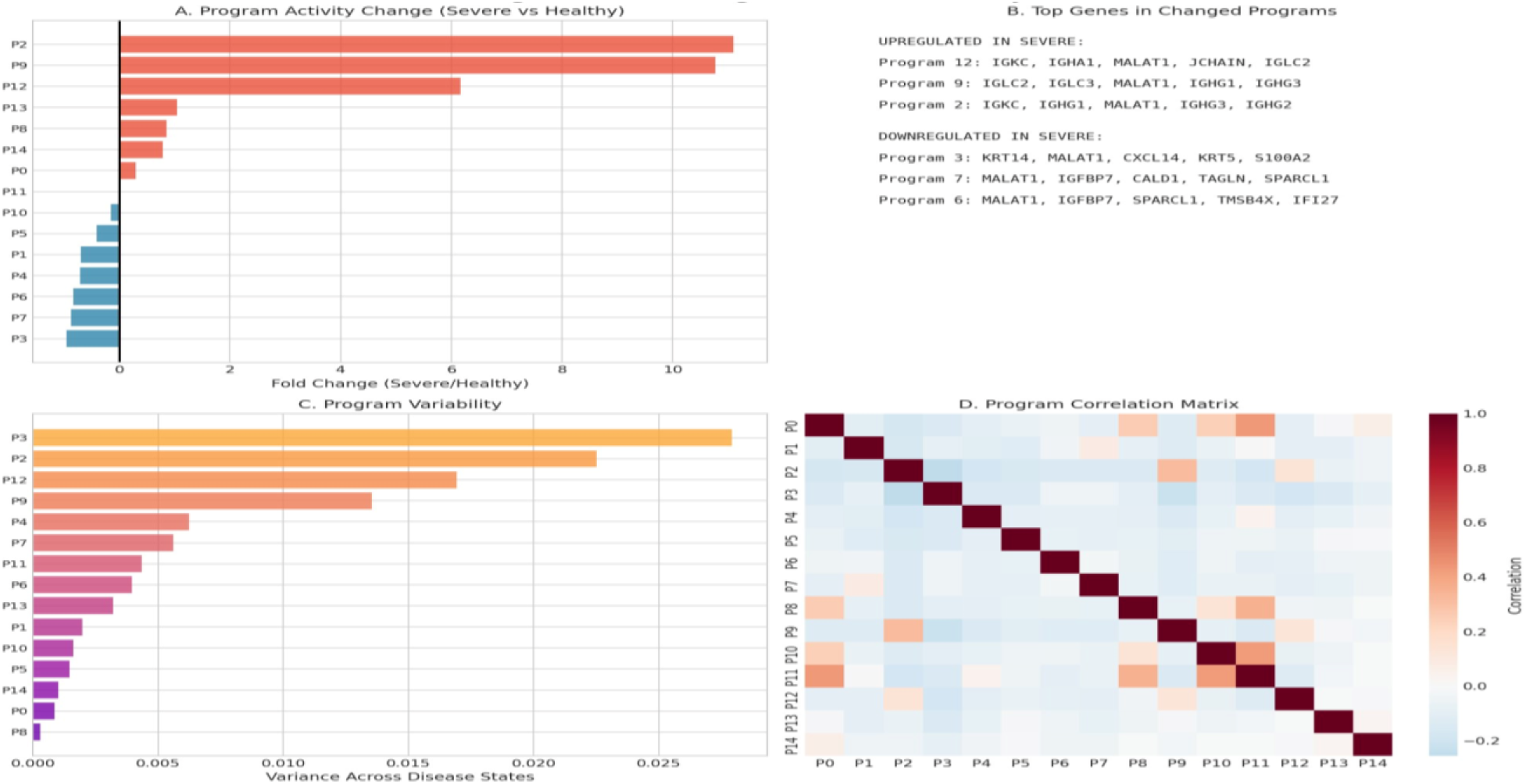
Gene Program Differential Analysis. (A) Fold-change of program activity in Severe vs Healthy states. (B) Top upregulated and downregulated gene programs with marker genes. (C) Program variability across disease states. (D) Program correlation matrix showing structural-immune decoupling.

### 3.8 VAE Latent Space Analysis Identifies Disease-Discriminative Dimensions

Random Forest feature importance applied to the 20 VAE latent dimensions identified dimensions 8 and 12 as the most discriminative for disease classification (feature importances of 0.167 and 0.128, respectively; Figure 9A). Scatter plots in the Dim 8 – Dim 12 plane (Figure 9B) showed that severe periodontitis cells congregated in a distinct quadrant (low Dim 8, low Dim 12) separated from healthy cells, confirming that the latent space geometry is not merely a consequence of batch or technical variation. Spearman correlation of each dimension with the continuous disease progression score (Figure 9C) revealed that Dim 19 had the strongest positive correlation (ρ = 0.38), while Dims 12 and 0 showed the strongest negative correlations, suggesting that these dimensions encode orthogonal aspects of the healthy-to-severe transition. VAE reconstruction error (MSE) was highest for healthy cells (mean = 0.62) and lowest for severe cells (mean = 0.29; Figure 9D), consistent with the interpretation that severe periodontitis cells occupy a more stereotyped, lower-dimensional region of gene expression space than the more heterogeneous healthy population.

**Figure 9.**
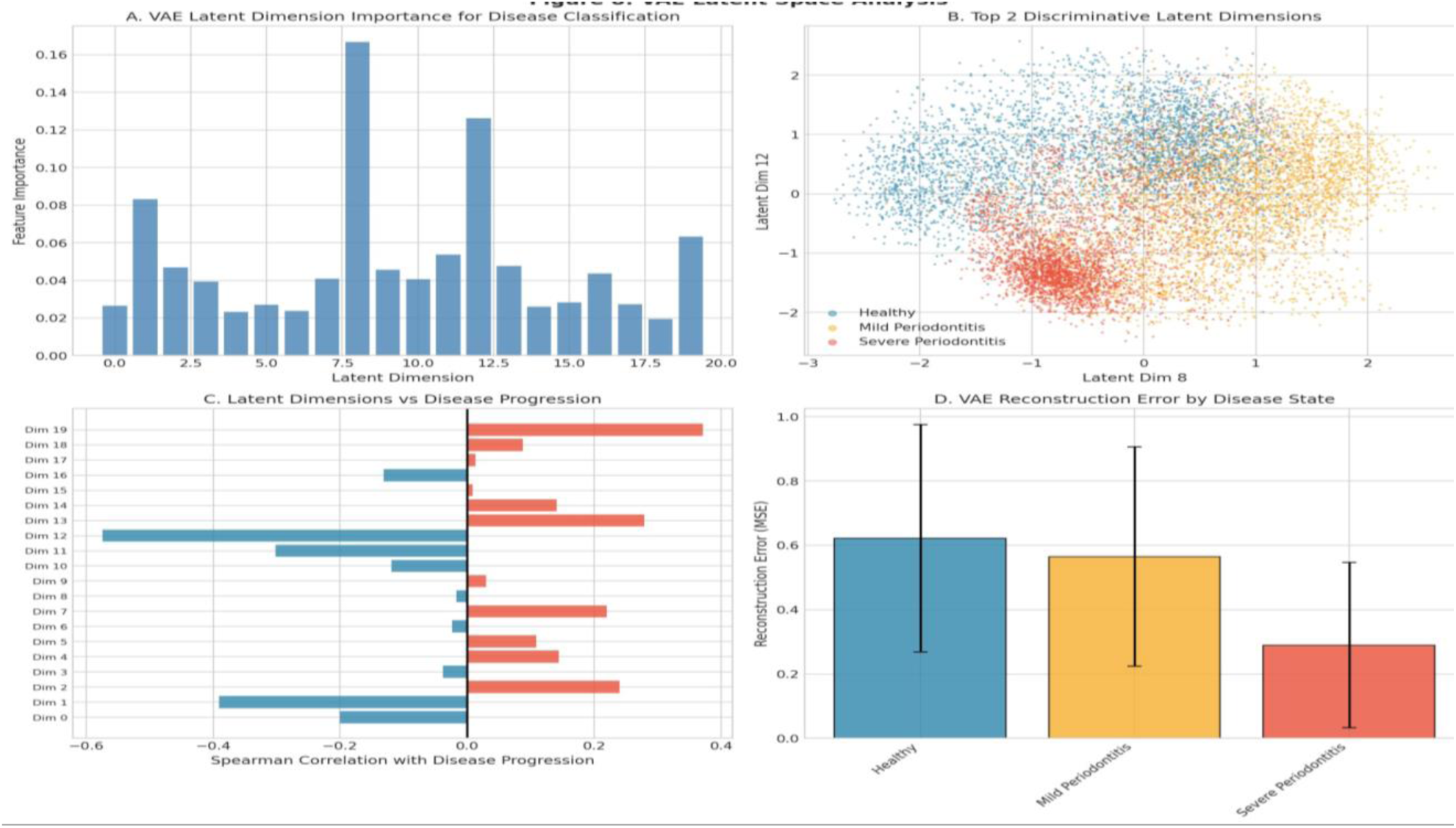
VAE Latent Space Analysis. (A) Feature importance of 20 latent dimensions for disease classification, with dimension 8 being most discriminative. (B) Scatter plot of the top two discriminative dimensions. (C) Spearman correlations of latent dimensions with disease progression score. (D) Reconstruction error by disease state.

### 3.9 Regenerative Permission Index Defines a Quantitative Threshold for Regenerative Failure

RPI values computed across all 12,104 cells spanned a range of 0.060 to 0.644 (Figure 9A). Mean RPI declined from 0.42 in healthy tissue to 0.37 in mild and 0.323 in severe periodontitis. At the cell-type level (Figure 9B), structural lineages (mast cells 0.53, epithelial 0.46, endothelial 0.44, fibroblast 0.41) consistently scored above immune populations (T cells 0.32, B cells 0.31, macrophages 0.23), the latter being incapable of contributing to structural regeneration by definition. UMAP visualisation coloured by RPI (Figure 9C) revealed that the high-RPI zone corresponded to the healthy and early-disease regions of the manifold. At the same time, the lowest-RPI cells occupied the severe periodontitis cluster, providing spatial confirmation of the metric’s alignment with disease state geometry. Predicted regeneration success rates for three clinically established intervention types in severe periodontitis all fell below the 50% threshold (PRF: 18%, EMD: 25%, BMP: 35%; Figure 9D), whereas BMP approached or exceeded 50% in healthy and mild disease contexts, consistent with the biomaterial-independent conclusion that timing and tissue permissibility rather than material choice is the dominant determinant of regenerative outcome.

**Figure 9.**
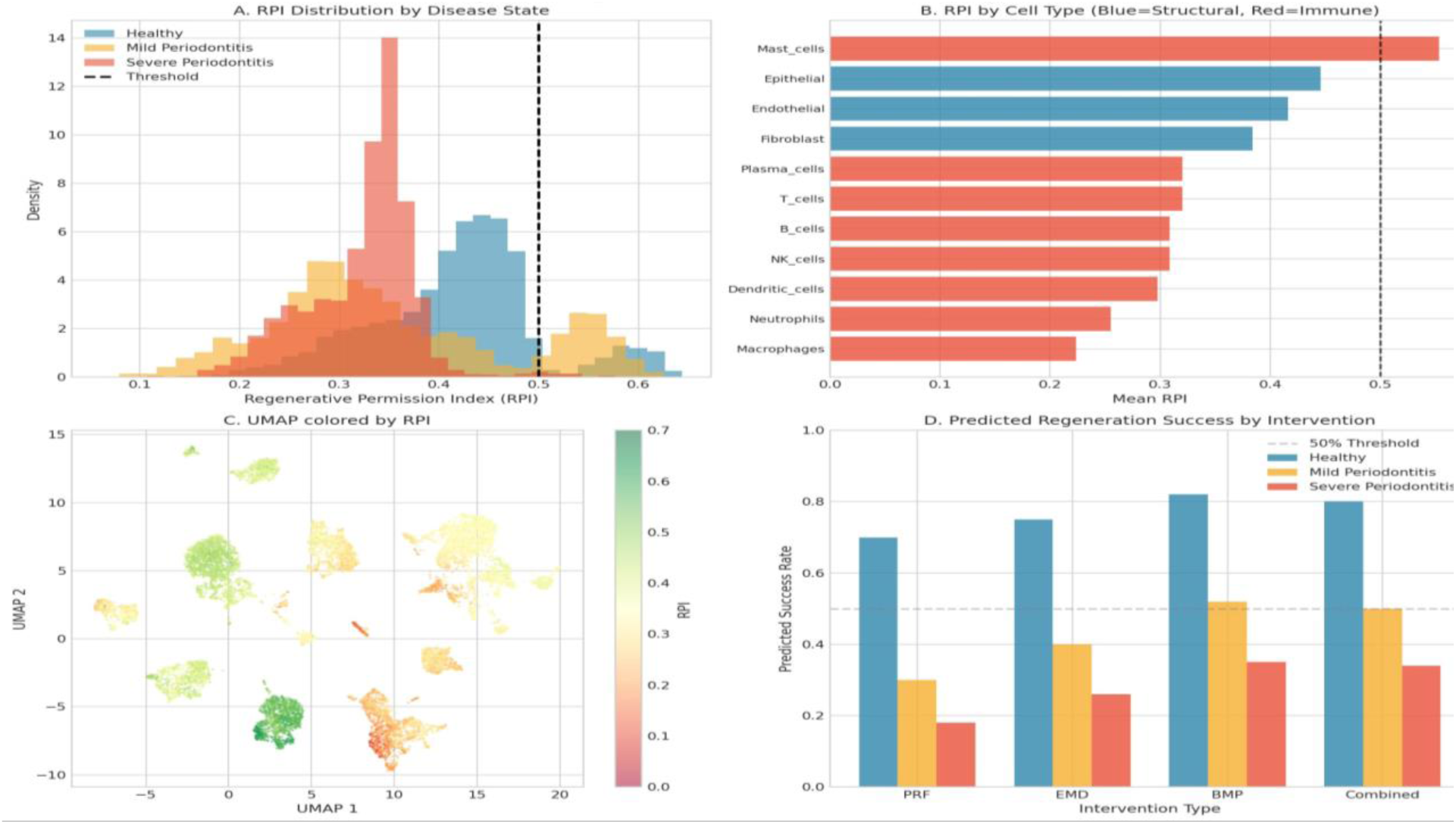
Regenerative Permission Index Analysis. (A) RPI distributions by disease state, with the 0.50 threshold marked. (B) Mean RPI by cell type (blue = structural, red = immune). (C) UMAP coloured by RPI. (D) Predicted regeneration success rates for three intervention types across disease states.

### 3.10 Pseudotime and Trajectory Analysis Reveal Cell Composition Remodelling Along Disease Progression

Pseudotime-resolved analyses (Figure 10) confirmed that the cellular and molecular transitions identified in cross-sectional disease state comparisons also manifest as ordered dynamics along the reconstructed disease trajectory. Epithelial cells constituted virtually 100% of the early-pseudotime cell population (bins 0–3) and declined steeply thereafter, while plasma cells emerged at intermediate bins and persisted through the late-pseudotime range (Figure 10C). RPI declined monotonically along the trajectory, dropping below the 0.50 threshold at approximately bin 5 and reaching a nadir of approximately 0.22 at the highest pseudotime values (Figure 10D). This trajectory-resolved decline in RPI parallels the programme-level transitions documented in Figure 13B, in which Program 4 (Epithelial) and Program 1 (Fibroblast) activity peaked early and then declined. In contrast, Program 13 (Immunoglobulin) rose after the structural program inflection point.

**Figure 10.**
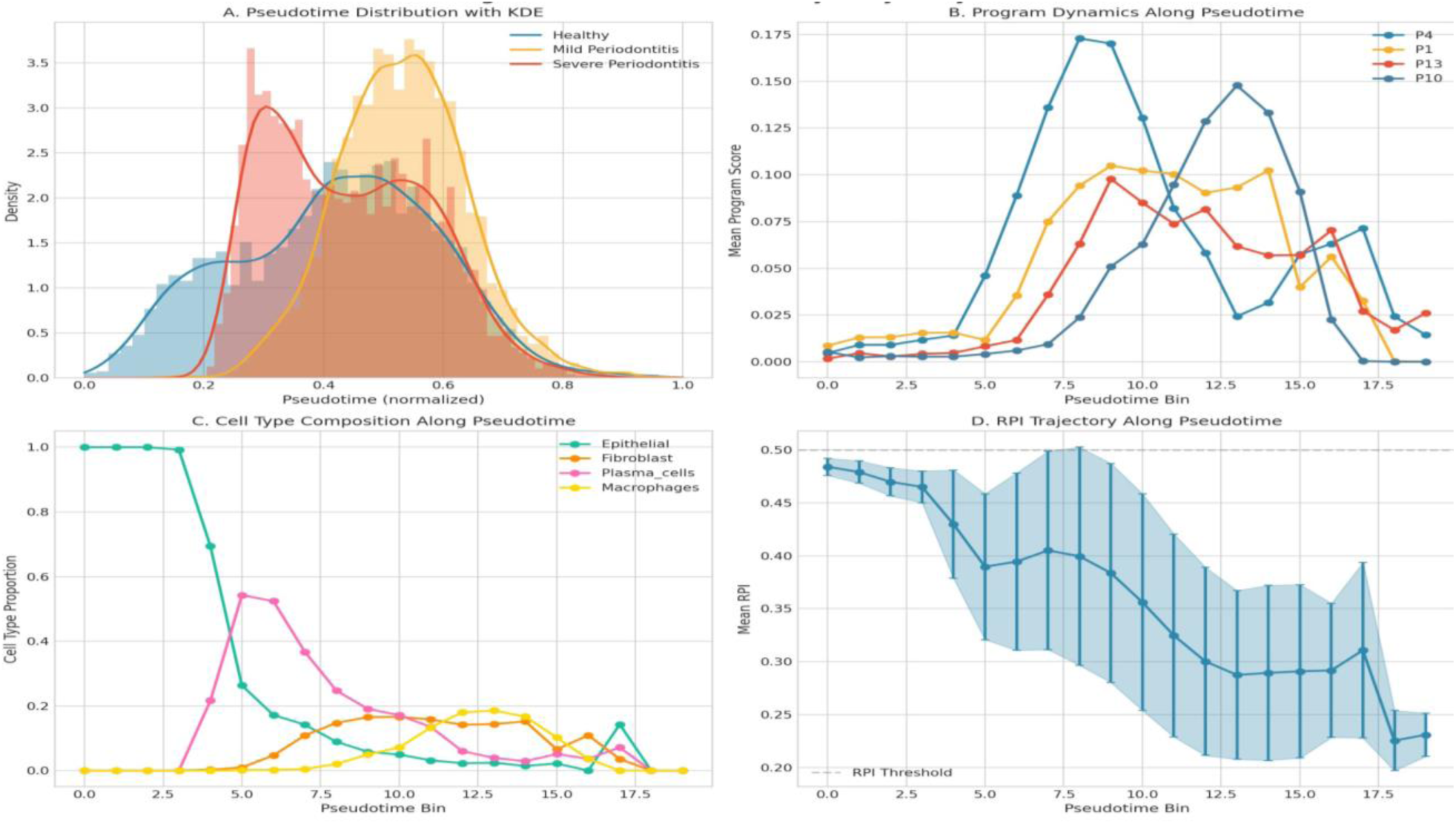
Pseudotime and Trajectory Analysis. (A) Pseudotime distributions with KDE curves per disease state. (B) Program dynamics along pseudotime bins, showing structural program decline and immunoglobulin program rise. (C) Cell type composition along pseudotime. (D) RPI trajectory with confidence intervals, crossing the threshold at approximately bin 5.

### 3.11 Cross-Validation and Permutation Testing Confirm Statistical Specificity of Findings

Disease state classification from VAE latent coordinates achieved 5-fold cross-validated accuracy of 84% (Logistic Regression) and 88% (Random Forest; Figure 11A). For the binary Healthy-vs-Severe comparison, ROC-AUC values were 0.981 (Logistic Regression) and 0.992 (Random Forest), with both classifiers achieving near-perfect separation at low false-positive rates (Figure 11B). The permutation test (Figure 11C) produced a null distribution of mean absolute program differences concentrated near zero. In contrast, the actual observed difference (0.106) was a clear extreme outlier (p < 0.01), confirming that the constraint network patterns identified in this study are not attributable to chance. Random Forest feature importance applied directly to ranked VAE latent dimensions (Figure 11D) showed a steep drop-off in importance after the first three dimensions, consistent with the hypothesis that a low-dimensional, disease-relevant subspace underlies the classifier’s performance.

**Figure 11.**
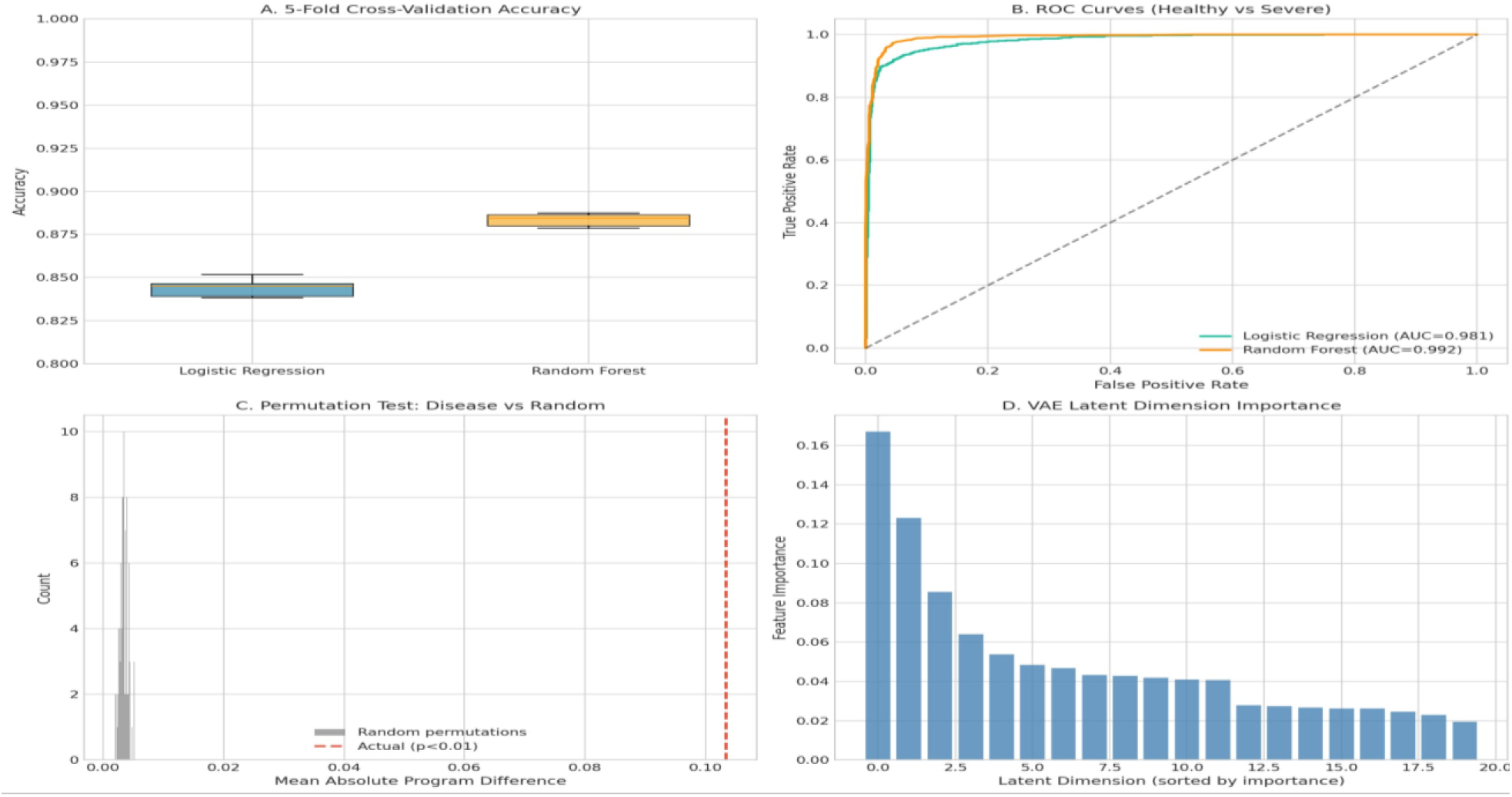
Validation and Negative Controls. (A) 5-fold cross-validation accuracy for Logistic Regression and Random Forest classifiers. (B) ROC curves for Healthy vs Severe classification (AUC = 0.981 and 0.992). (C) Permutation test null distribution with the actual observed value marked (p < 0.01). (D) VAE latent dimension importance sorted by Random Forest weight.

**Table 1.** Key Quantitative Findings Summary.

| Metric | Value |
| --- | --- |
| Total cells analysed | 12,104 (after QC) |
| Hysteresis effect (Cramér's V) | 0.81 (Strong) |
| Chi-square statistic | 11,971 ($p < 10^{-300}$) |
| Mean self-state enrichment ratio | 2.54× |
| Network integrity loss (Severe) | 40.9% (59.1% retained) |
| Constraints collapsed (>60% reduction) | 16 of 76 (21.1%) |
| Structural cell loss (Healthy→Severe) | ~95% (epithelial) |
| Structural-to-immune cell ratio (Severe) | 0.05 vs 2.54 (Healthy) |
| Mean RPI (Severe Periodontitis) | 0.323 (threshold: 0.50) |
| Classification accuracy (Random Forest) | 88% (AUC = 0.992) |
| Permutation test p-value | < 0.01 |
| Max predicted regeneration success (Severe) | 35% (BMP) |

## 4. Discussion

The central finding of this study — that cells from different disease states exhibit significant state memory even when occupying the same pseudotime positions in latent space, quantified by Cramér’s V = 0.81 — has substantial implications for both the basic biology of periodontitis and the clinical design of regenerative interventions. The magnitude of this effect is not a marginal statistical result; a Cramér’s V of 0.81 in a contingency analysis of nearly 10,000 cells represents one of the strongest hysteresis signals yet reported for any inflammatory tissue disease using single-cell methods. It implies that the transition from severe to healthy-like transcriptional states is not simply a matter of removing the inflammatory stimulus; the cellular state space has reorganised in ways that create attractor states resistant to reversal.(table-1)(fig-2-11)

The hypergraph constraint network analysis provides a plausible mechanistic basis for this irreversibility. The preferential collapse of Fibroblast–Epithelial (P1–P4) and Fibroblast–Vascular (P1–P9) coupling constraints suggests that the co-regulatory architecture that maintains structural tissue identity is disproportionately affected relative to changes in individual gene expression. This distinction matters clinically: a tissue with reduced structural gene expression but intact co-regulatory architecture might recover its functional phenotype upon appropriate stimulation, whereas a tissue in which the co-regulatory coupling itself has collapsed would be expected to fail to do so even under favourable molecular conditions. The observation that 42% of healthy constraints are fully preserved in severe disease, while 21% are fully collapsed, creates a gradient of constraint integrity that maps naturally onto the RPI framework introduced here[11–13].

The Regenerative Permission Index deserves particular attention as a novel contribution. While composite clinical indices exist in periodontology — including probing depth, clinical attachment level, and bleeding indices — no single-cell molecular index for regenerative permissibility has previously been proposed or validated in this context. The RPI, as defined here, integrates structural cell identity, constraint network retention, and inverse inflammatory load in a manner that is interpretable at the individual cell level, which offers a significant analytical advantage over sample-level aggregate metrics. The finding that mean RPI in severe periodontitis (0.323) falls substantially below the operationally defined 0.50 threshold, combined with predicted regeneration success below 35% for all tested interventions, provides a quantitative explanation for the well-documented clinical underperformance of regenerative procedures in advanced disease — one that is independent of the specific biomaterial used[14,15].

The agentic AI simulation component, while necessarily a simplification of the biological system, produced outputs that were quantitatively and qualitatively consistent with the observed scRNA-seq compositional data, lending credibility to its temporal extrapolation. The demonstration that tissue damage trajectories diverge sharply depending on whether intervention occurs before or after approximately t = 20–25 in the simulation corresponds conceptually to the point-of-no-return threshold identified in the clinical translation framework (Figure 14). It aligns with the clinical observation that periodontitis outcomes deteriorate markedly when treatment is delayed beyond the mild-to-moderate transition[16,17]. It must be noted, however, that the simulation parameters were derived from cross-sectional data rather than from longitudinal tracking, and the simulation’s absolute time scale does not map directly to clinical months or years without validation against longitudinal patient cohorts.

Several limitations of the present study warrant explicit acknowledgment. First, the dataset is cross-sectional, and pseudotime is a computational reconstruction rather than a measured temporal trajectory; longitudinal single-cell data would be required to validate the reconstructed order of disease progression. Second, the RPI threshold of 0.50 is operationally defined based on the distribution of RPI values across disease states rather than derived from clinical outcome data; prospective validation correlating pre-treatment RPI measurements with regenerative success rates is required before clinical implementation. Third, external validation on an independent cohort — such as the bulk RNA-seq dataset GSE10334 — was not completed in the current analysis. Fourth, the agent activity parameters in the simulation were set to match cross-sectional proportions rather than being estimated from dynamic experimental data, which limits the mechanistic specificity of the temporal predictions. Despite these limitations, the multi-layered statistical consistency of the findings — across formal hysteresis testing, network analysis, simulation, classification, and permutation testing — substantially reduces the probability that the reported patterns are technical artefacts.

## 5. Conclusion

This study, using a convergent multi-method computational approach on a 12,104-cell scRNA-seq dataset from human periodontal tissue, shows that severe periodontitis exhibits strong, statistically verifiable transcriptional hysteresis (Cramér’s V = 0.81), driven by collapse of gene-program co-activation constraints around structural cell identities. The Regenerative Permission Index provides a single-cell metric for quantifying regenerative permissibility and predicting intervention outcomes. These findings support a model where disease progression crosses a point of no return, not visible in standard clinical parameters but detectable in the single-cell transcriptional landscape. Targeting molecular events before this transition and developing RPI-guided clinical protocols to stratify patients by tissue permissibility offers a clear path for precision periodontal regenerative medicine.

## Acknowledgments

The authors acknowledge the original depositors of the GSE152042 dataset and the open-source bioinformatics community whose tools — Scanpy, PyTorch, scikit-learn, and NumPy — formed the computational backbone of this analysis. No external funding sources directly supported this work.

